# SKIDA1 transiently sustains MLL::ENL-Expressing hematopoietic progenitors during neonatal stages and promotes B-lineage priming

**DOI:** 10.1101/2025.04.12.648529

**Authors:** Jonny Mendoza-Castrejon, Wei Yang, Elisabeth Denby, Rohini Muthukumar, Emily B. Casey, Riddhi M. Patel, Sarah K. Tasian, Jeffrey A. Magee

## Abstract

Infant leukemias arise as B-cell acute lymphocytic (B-ALL) or acute myeloid leukemia (AML). The majority are driven by chromosomal rearrangements of the *MLL/ KMT2A* gene (*MLL*r) and arise in utero, implying a fetal cell of origin. Fetal and neonatal hematopoietic progenitors have unique transcriptomes and epigenomes, raising the question of whether MLL fusion proteins activate distinct target gene profiles during these early stages of life. Here, we use a transgenic mouse model of MLL::ENL-driven leukemia to identify *Skida1* as a target gene that is more highly induced in fetal and neonatal progenitors than in adult progenitors. *SKIDA1* is highly expressed in human *MLL*r leukemias and the protein associates with the Polycomb Repressive Complex 2 (PRC2). We show that *Skida1* is dispensable for normal hematopoiesis, but it promotes B-cell priming and maintains MLL::ENL-expressing hematopoietic stem cells (HSCs) and multipotent progenitor cells (MPPs) during neonatal development. Conditional deletion of *Skida1* has no effect on normal HSC function, yet it impairs B-cell production from neonatal MLL::ENL-expressing HSCs while leaving myeloid leukemogenesis unaffected. Temporally-restricted targets of MLL fusion proteins, such as SKIDA1, can therefore tune cell fates at different ages, potentially influencing the types MLLr leukemias that arise at different ages.

## INTRODUCTION

Infant leukemias can present as either acute myeloid leukemia (AML) or B-cell acute lymphoblastic leukemia (B-ALL).^1^ A majority of infant leukemias are driven by chromosomal rearrangements that involve the *MLL* (*KMT2A*) gene. The rearranged loci express MLL fusion proteins (e.g., MLL::ENL, MLL::AF9 and MLL::AF4) that ectopically activate *HOX* genes and *MEIS1* to drive leukemic transformation.^2–5^ *MLL* rearrangements (MLLr) have been shown to arise before birth in patients who ultimately developed infant leukemias, indicating a fetal/neonatal cell of origin.^6–9^ This raises the question of whether MLLr activate distinct leukemogenic programs when they arise in fetal/neonatal progenitors as compared to adult progenitors. Fetal- or neonatal-specific effectors of transformation might explain why infant leukemias are often less responsive to chemotherapy than non-infant childhood or adult leukemias.

Prior studies have shown that fetal and neonatal progenitors do have distinct transcriptional and functional responses to MLL fusion proteins. For example, we previously used a transgenic mouse model to test whether MLL::ENL induces AML with varying degrees of efficiency across fetal, neonatal and adult stages of development.^10^ Neonatal MLL::ENL induction initiated AML with greater efficiency than fetal or adult MLL::ENL induction, suggesting that transformation efficiency peaks shortly after birth. Likewise, MLL::AF9 and MLL::AF4 have been shown to transform human umbilical cord progenitors more efficiently than adult bone marrow progenitors.^11,12^ The data suggest that MLL fusion proteins activate subsets of target genes more effectively in fetal and neonatal progenitors than in adult progenitors. Such targets might potentiate transformation, or they might alter lineage biases within pre-leukemic cells of origin.

To understand how age alters transcriptional responses to MLL fusion proteins, we used a previously described, inducible model of MLL::ENL-driven AML to identify target genes that distinguish fetal, neonatal and adult progenitors.^10,13^ We identified *Skida1* as a gene that is more highly induced in fetal and neonatal progenitors than in adult progenitors. *SKIDA1* is highly expressed in human MLLr leukemias, and the encoded protein associates with several components of the Polycomb Repressive Complex 2 (PRC2). We generated germline and conditional *Skida1* loss of function mouse models and found that it is dispensable for normal murine hematopoiesis. In contrast, *Skida1* sustains MLL::ENL-expressing pre-leukemic progenitors during neonatal stages of development, particularly HSCs, and it selectively maintains B-cell potential. *Skida1* deficiency did not impede myeloid leukemogenesis, and *SKIDA1* is not necessary to sustain fully transformed infant B-ALL. Altogether, the data show that *Skida1* is an effector of MLL::ENL-driven B-lineage priming and HSC/MPP maintenance during neonatal stages of life, prior to transformation, but it is not required to sustain infant leukemias after transformation.

## RESULTS

### *Skida1* is an MLL::ENL target gene that is more highly expressed in fetal and neonatal progenitors than in adult progenitors

To identify genes that are more strongly induced by MLL::ENL in fetal and neonatal progenitors than in adult progenitors, we analyzed MLL::ENL-driven changes in gene expression in HSC/MPPs (Lineage-Sca1+Kit+; LSK) at embryonic day (E)16, postnatal day (P)14 and 8 weeks after birth. For these experiments, we used mice with *Vav1-Cre, Rosa26^LoxP-STOP-LoxP-reverse-tet-transactivator^*(*Rosa26^LSL-rtTA^*) and *Col1a1^TetO-MLL::ENL^* alleles (hereafter referred to as Tet-On-ME). These mice express MLL::ENL specifically in hematopoietic cells upon exposure to DOX. We cultured LSK cells for 48hrs, with or without DOX, and then performed RNA-sequencing (RNA-seq) to identify changes in gene expression that occurred shortly after MLL::ENL induction, independently of microenvironmental differences that might distinguish fetal, neonatal and adult progenitors.

MLL::ENL induced extensive changes in gene expression for each age studied (Figure 1A, Table S1). A majority of differentially expressed genes were induced or repressed to similar levels in fetal, neonatal and adult progenitors (Figure 1A, Table S1). However, several genes had fetal- or neonatal-biased patterns of expression (Figure 1A, highlighted cluster, Table S2), including the known MLL::ENL target gene *Hoxa9* (Figure 1A). In addition, B-cell identity genes, including *Rag1*, *Rag2*, *Bach2* and *Bank1*, were more highly expressed in E16 and P14 progenitors (Figure 1A-C), indicating that MLL::ENL enhances B-lymphoid priming in fetal/neonatal progenitors. Several Hallmark gene signatures, including MYC target genes, Oxidative Phosphorylation genes and E2F target genes, were downregulated in MLL::ENL expressing progenitors at all ages based on Gene Set Enrichment Analysis (GSEA) (Figure 1D). Interferon and TGF-beta signatures were activated by MLL::ENL specifically in P14 and adult progenitors (Figure 1D). These observations reinforce the premise that MLL::ENL induces distinct transcriptional programs in fetal, neonatal and adult progenitors, independent from any microenvironmental differences.

**Figure 1.**
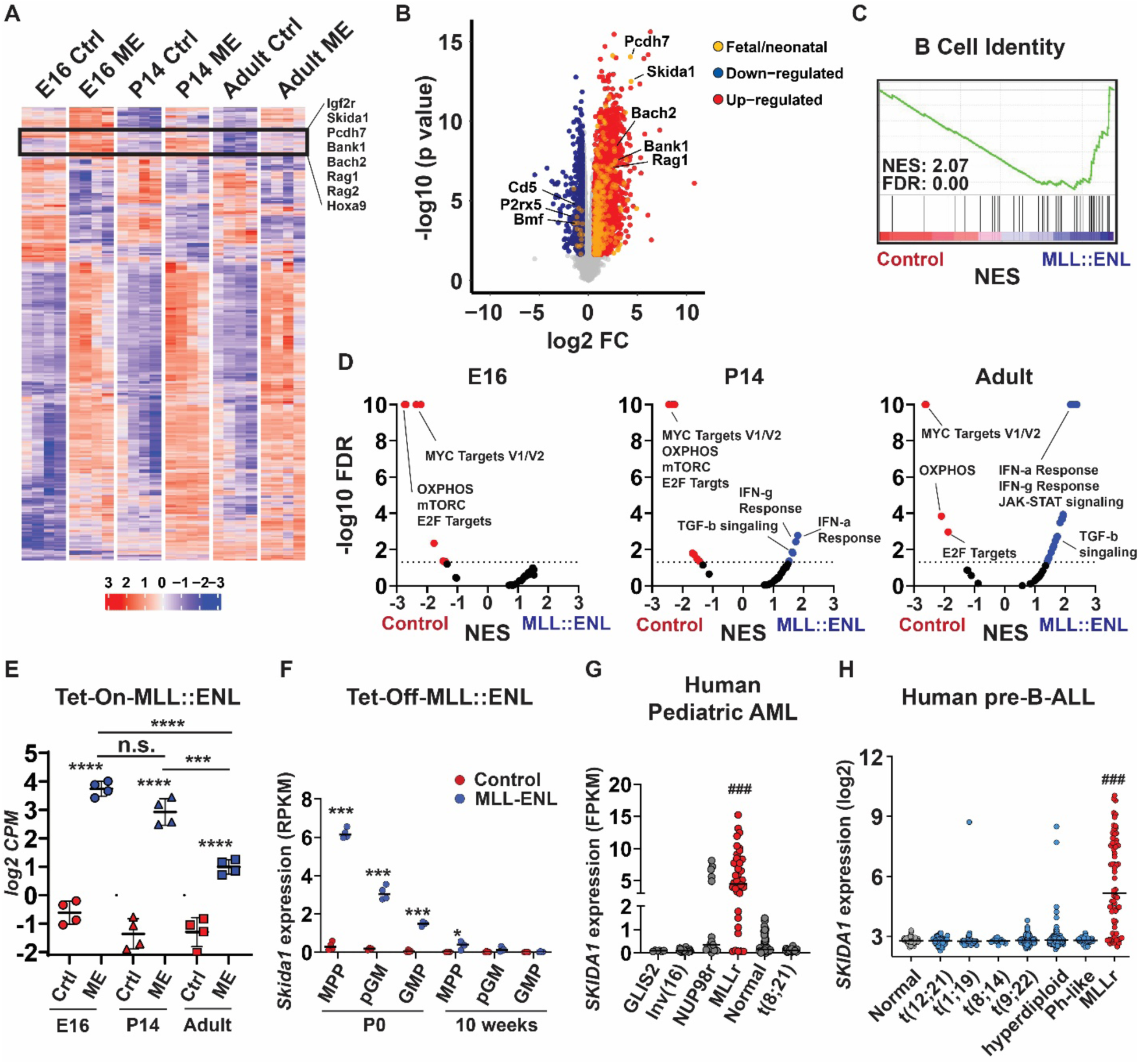
MLL::ENL induces *Skida1* expression in fetal and neonatal HSC/MPPs. (A) Heatmap showing genes that are significantly differentially expressed (FDR<0.05, log2(fold change) > 0.6) upon MLL::ENL induction in E16, P14 and 8-week-old adult Tet-On-ME LSK cells. A cluster of genes that are more highly induced in fetal and neonatal LSK cells, relative to adult LSK cells, is outlined with representative genes indicated. (B) Volcano plot indicating genes that are significantly differentially expressed in fetal LSK upon MLL::ENL induction, with color coding to indicate genes that are selectively induced in fetal and neonatal progenitors. (C) GSEA plot showing B-cell signature enrichment in E16 MLL::ENL-expressing progenitors. (D) Volcano plots indicating changes in expression in Hallmark gene signatures upon MLL::ENL expression, based on GSEA. NES indicates normalized enrichment scores. (E) *Skida1* expression in LSK cells from the indicated ages, based on RNA-seq. n= 4 biological replicates per genotype and age. ***p<0.001, ****p<0.0001 by one-way ANOVA with Tukey’s posthoc test. (F) *Skida1* expression in wild-type and Tet-Off-ME progenitors at P0 and 10-weeks-old based on previously described RNA-seq data.^10^ n= 4 biological replicates per progenitor population and age. *p<0.05, ***p<0.001 by Student’s t-test. (G, H) *SKIDA1* expression in MLLr pediatric AML and pre-B-ALL collected under the auspices of TARGET (www.cancer.gov/ccg/research/genome-sequencing/target) and TCGA (www.cancer.gov/tcga) sequencing projects, respectively. ###p<0.001 relative to all other subtypes by one-way ANOVA with Tukey’s posthoc test.

The list of genes that were more highly induced by MLL::ENL in fetal/neonatal HSC/MPP, as compared to adult HSC/MPP, included *Skida1*, a gene that lacks a previously-described role in infant or MLLr leukemogenesis (Figure 1E). Analysis of previously collected *in vivo* RNA-seq data^10^ showed that *Skida1* is more highly expressed *in vivo* in P0 MLL::ENL-expressing HSC/MPPs, relative to P14 and 8-weeks-old HSC/MPPs (Figure 1F). Fetal pre-granulocyte-monocyte progenitors (pGM) and granulocyte-monocyte progenitors (GMPs) showed somewhat lower levels of *Skida1* induction by MLL::ENL at P0 (Figure 1F). *Skida1* expression was very low in all normal cell populations at all ages. Thus, *Skida1* is a fetal/neonatal MLL::ENL target gene both in *ex vivo* culture experiments and *in vivo*. Previously collected RNA-seq data representing human pediatric AML or human pre-B-ALL showed high *SKIDA1* expression almost exclusively in MLLr leukemias (Figure 1G, H). These data raise questions as to how SKIDA1 functions at a molecular level and whether it promotes MLLr leukemogenesis in fetal/neonatal progenitors.

### SKIDA1 is a nuclear protein that interacts with the Polycomb Repressive Complex 2

*Skida1* encodes a 98 kDa protein with an N-terminal SKI/DACH domain and a C-terminal EPOP (Elongin BC and Polycomb Repressive Complex 2-associated Protein) homology region (Figure 2A). A SKIDA1-Green Fluorescent Protein (GFP) fusion localized primarily to the nucleus (Figure 2B, C), suggesting potential interactions with other nuclear proteins and transcriptional regulators. Indeed, SKI/DACH domains have been shown to bind SMAD family transcription factors, and EPOP has been shown to interact with PRC2.^14–20^ These features suggest that SKIDA1 may interact with either SMAD family proteins or PRC2.

**Figure 2.**
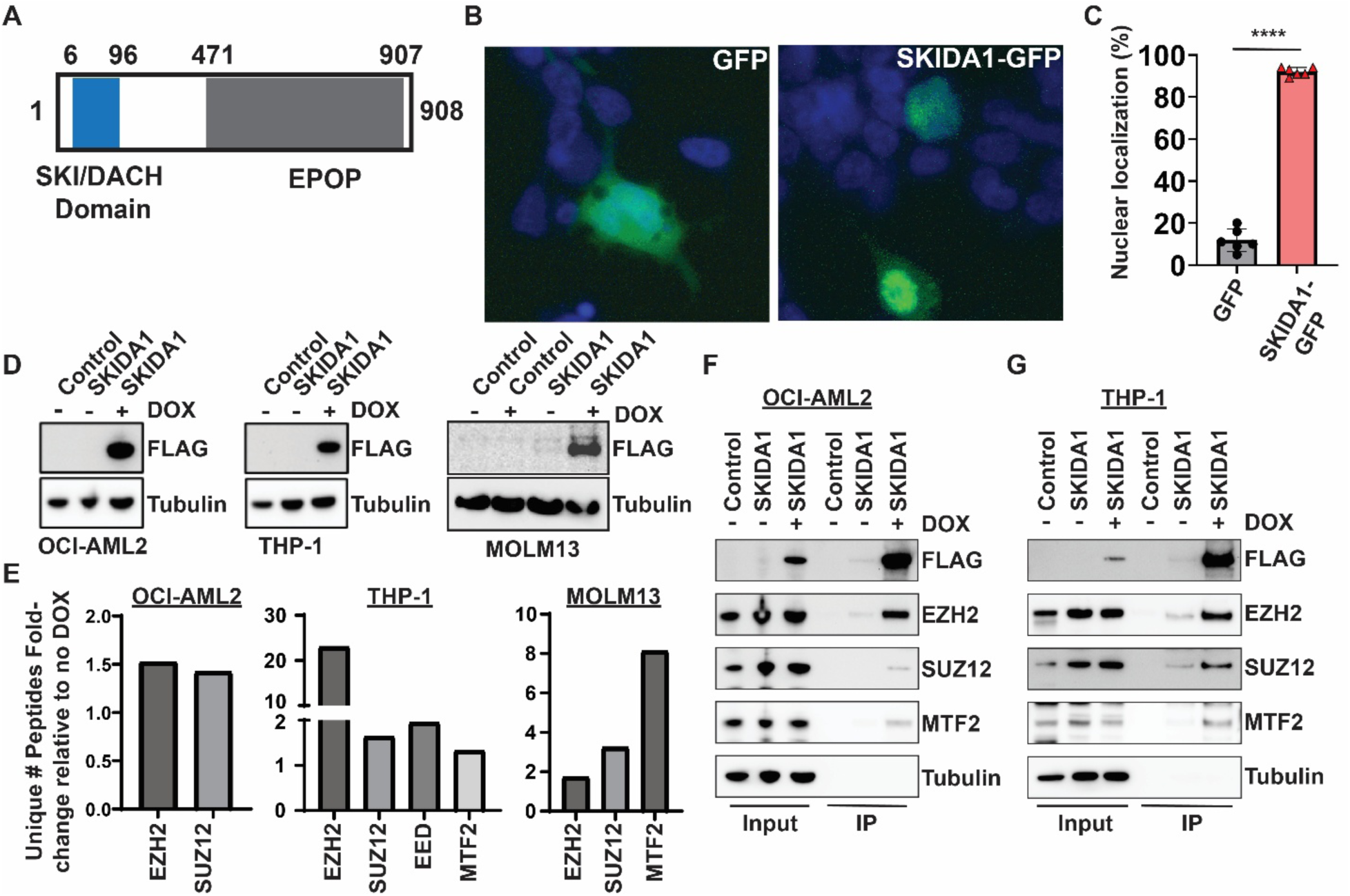
SKIDA1 is a nuclear protein that interacts with PRC2 proteins. (A) Schematic of the SKIDA1 protein highlighting the SKI/DACH (blue) and EPOP homology (gray) domains with corresponding human amino acid numbers. (B) Representative pictures of GFP (left) and SKIDA1-GFP (right) subcellular localization in transfected 293T cells. (C) Quantification of cells with nuclear SKIDA1-GFP localization. ****p<0.0001 by Student’s t-test. (D) Western blot showing FLAG-SKIDA1 expression in in human OCI-AML2, THP-1 and MOLM13 cell lines after 3 days of DOX exposure. (E) Fold increase in unique peptides identified in FLAG-SKIDA1 pulldown assays by LC-MS/MS in OCI-AML2, THP-1 and MOLM13 cell lines, based on no DOX vs. DOX comparison. (F, G) Co-immunoprecipitation assays and Western blots confirming interactions between FLAG-SKIDA1 and the indicated PRC2 proteins in OCI- AML2 and THP-1 cells.

To test whether SKIDA1 interacts with PRC2 or other nuclear proteins, we performed co-immunoprecipitation assays followed by mass spectrometry. We generated a DOX-inducible SKIDA1-3xFLAG lentiviral construct (SKIDA1-FLAG) and transduced THP-1, OCI-AML2 and MOLM13 AML cells to generate stable, inducible MLLr cell lines (Figure 2D). We immunoprecipitated FLAG-SKIDA1 from DOX-treated and untreated cells and performed on-bead protein digestion followed by liquid chromatography – mass spectrometry (LC-MS/MS). We identified several PRC2 proteins, including EZH2, SUZ12, EED and MTF2, as having >1.5-fold higher peptide counts in DOX-treated cells, relative to untreated cells across the three cell lines (Figure 2E; Table S3). PRC2 associated proteins were also enriched in a similar experiment performed with 32D cells (Table S4), an immortalized mouse progenitor cell line. Co-immunoprecipitation assays validated the strong interaction between SKIDA1 and EZH2 (Figure 2F, G), as well as weaker interactions with SUZ12 and MTF2. Altogether, these data show that SKIDA1 is a nuclear protein that interacts with the PRC2 complex. Prior work has shown that PRC2 regulates normal fetal/neonatal hematopoiesis and that the complex is necessary to sustain leukemogenesis.^21–26^ These observations raise the question of whether *Skida1* regulates hematopoiesis and leukemogenesis.

### *Skida1* is dispensable for normal murine hematopoiesis and HSC function

To test whether *Skida1* regulates normal development, we generated mice with a germline *Skida1* loss-of-function allele, and we analyzed the mice for morphologic abnormalities and hematopoietic defects. We used CRISPR/Cas9 to delete the single *Skida1* coding exon to generate a null allele (Figure 3A, B). *Skida1^-/-^* mice were born at normal Mendelian ratios but grew less than wildtype or *Skida1^+/-^* littermates over the first 21 days of life (Figure 3C, D). Many *Skida1*-deficient mice did not survive past weaning (Figure S1A), and those that did survive remained small into adulthood (Figure 3D). The most obvious gross morphologic defect in *Skida1^-/-^*mice was failure to develop normal eyes. At birth, *Skida1^-/-^* mice had partially opened eyelids, in contrast to wildtype littermates (Figure 3E, left panel). By adulthood, the orbits were scarred shut and the eyes decayed (Figure 3E, right panel). Thus, in addition to being an MLL::ENL target gene, *Skida1* regulates eye development.

**Figure 3.**
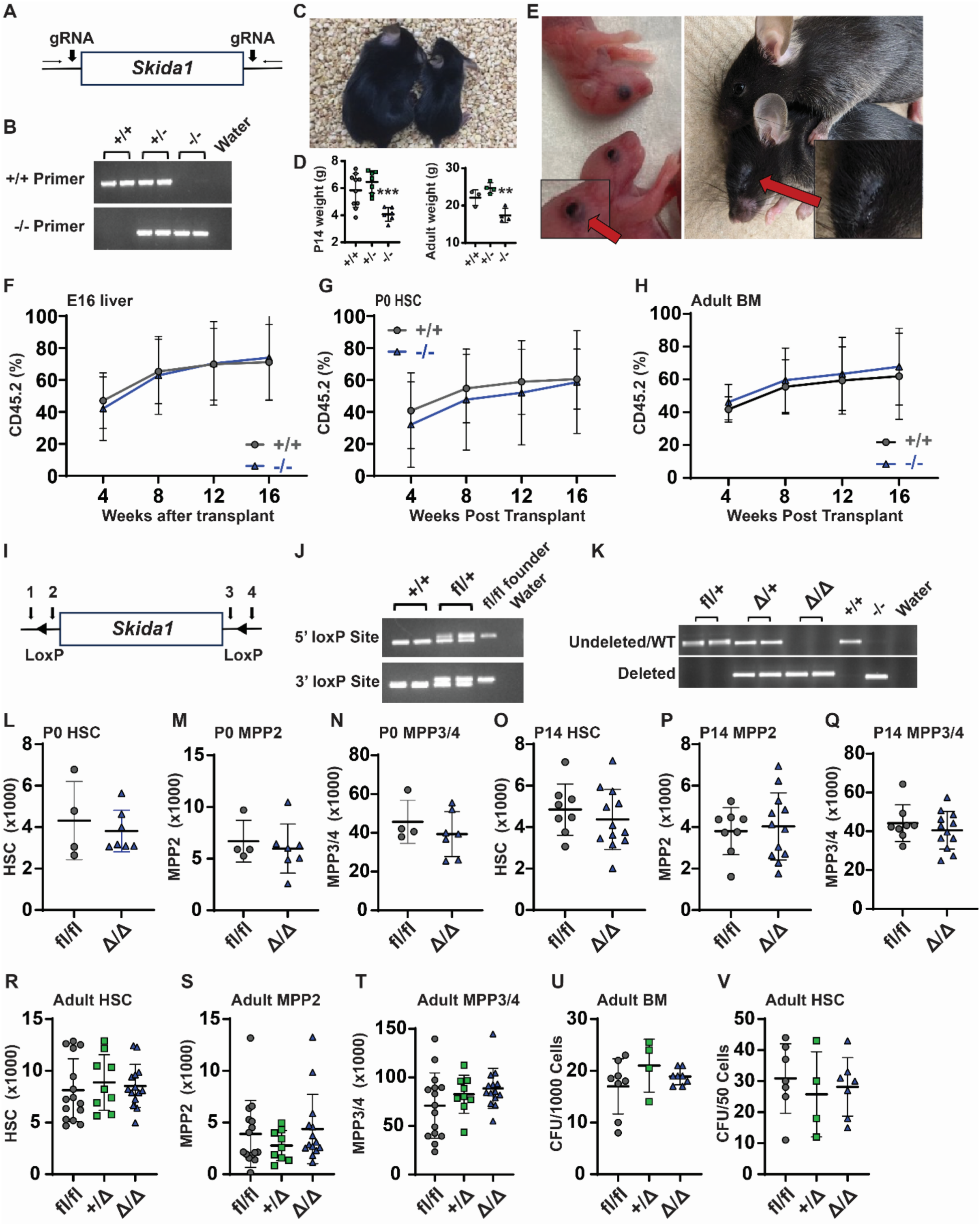
*Skida1* is not required for normal hematopoiesis or HSC function during fetal, neonatal, or adult stages. (A) Diagram of *Skida1* germline loss-of-function allele with vertical arrows indicating gRNA cut sites around the single coding exon to generate *Skida1* null mice. (B) Representative genotyping PCR of the deleted *Skida1* allele. (C) Representative pictures of P21 *Skida1*^-/-^ (right) and *Skida1^+/+^*littermates (left). (D) Body weights for *Skida1^+/+^*, *Skida1^+/-^* and *Skida1^-/-^* mice at P14 and 8 weeks old. P14 n=10 (+/+), n=7 (+/-), n=6 (-/-); 8 week n=3 (+/+), n=4 (+/-), n=4 (-/-). **p<0.01, ***p<0.001 by one-way ANOVA with Tukey’s posthoc test. (E) Representative pictures of eye defects in *Skida1*^-/-^ mice at P0 and 8 weeks old. (F-H) CD45.2+ peripheral blood chimerism levels in recipients of E16 fetal liver cells, P0 HSCs or 8-week-old bone marrow cells from donors with the indicated genotypes. E16 recipients n=17 (+/+), n=18 (-/-); P0 recipients n=17 (+/+), n=19 (-/-); and 8 week recipients n=9 (+/+), n=8 (-/-). (I) Diagram of *Skida1* conditional loss-of-function allele. LoxP sites were inserted around the single coding exon by Cas9 editing and homologous recombination. Genotyping primer sites (1-4) are indicated. (J) Representative genotyping PCR for the *Skida1^fl^* allele. (K) Representative PCR indicating complete deletion of the floxed allele in CD45.2^+^ peripheral blood cells from *Skida1^fl/fl^*; *Vav*-Cre mice. (L-T) Total cell numbers of indicated populations and genotypes in P0 whole liver, or P14 or adult bone marrow from two hind limbs (tibia and femur). P0 n=4 (fl/fl), n=7 (Δ/Δ); P14 n=8 (fl/fl), n=12 (Δ/Δ); Adult n=15 (fl/fl), n=9 (+/Δ), n=14 (Δ/Δ). (U, V) CFU frequencies generated from 1000 whole bone marrow cells or sorted HSCs from adult mice with the indicated genotypes. Whole bone marrow n=8 (fl/fl), n=4 (+/Δ), n=8 (Δ/Δ); HSCs n=7 (fl/fl), n=4 (+/Δ), n=7 (Δ/Δ). None of the comparisons in panels L-V indicated significant differences among the genotypes, as calculated by Student’s t-test or one-way ANOVA with Tukey’s posthoc test.

We analyzed hematopoiesis in E16, P0 and adult wildtype *Skida1^+/-^* and *Skida1^-/-^* mice. HSC (CD150+CD48-LSK) and MPP (CD48+LSK) numbers were similar across all 3 genotypes at E16 (Figure S1B, C). At P0, we observed a small increase in total HSC numbers in *Skida1^-/-^* pups but no differences in MPP numbers (Figure S1D, E).

Surviving adult *Skida1^-/-^* mice had similar numbers of HSCs and MPPs (Figure S1F, G). To assess HSC function, we competitively transplanted 300,000 fetal liver cells from E16 wildtype or *Skida1^-/-^* mice (CD45.2+) along with 300,000 wildtype competitor bone marrow cells (CD45.1+) into lethally irradiated CD45.1 recipient mice. We did not observe differences in donor peripheral blood chimerism over 16 weeks of post-transplant evaluation (Figure 3F, Figure S1H). We also transplanted 20 purified HSCs per recipient from P0 wildtype or *Skida1^-/-^* mice, along with 300,000 wildtype competitor cells, followed by secondary transplants of 3 million unfractionated primary recipient bone marrow cells. These assays did not reveal genotype-specific differences in repopulation in either primary or secondary recipients (Figure 3G, Figure S1I, K, L). Finally, we competitively transplanted 300,000 bone marrow cells from adult wildtype and *Skida1^-/-^* mice and again saw no difference in repopulating activity (Figure 3H, Figure S1J). Together, these data show that *Skida1* does not regulate normal fetal, neonatal or adult hematopoietic progenitor frequencies or HSC function, consistent with its low level of expression in the absence of MLL::ENL (Figure 1F).

The growth and survival defects of *Skida1^-/-^* mice confounded plans to evaluate pre-leukemic progenitors and leukemogenesis in the context of MLL::ENL expression so we generated a conditional loss-of-function *Skida1^flox^*allele. LoxP sites were inserted to flank the single *Skida1* exon (Figure 3I, J). We then used *Vav1*-*Cre* to delete *Skida1* in hematopoietic progenitors beginning at E10.5 (Figure 3K). Adult *Skida1^fl/fl^; Vav1-Cre* (*Skida1^Δ/Δ^*) mice had normal weights, bone marrow cellularity, spleen weights and peripheral white blood cell counts (Figure S1M-P). Furthermore, *Skida1* deletion did not alter HSC, MPP2 (CD150^+^CD48^+^LSK), MPP3/4 (CD150^-^CD48^+^LSK) or granulocyte-monocyte progenitor (Lineage^-^Sca1^-^Kit^+^CD105^-^CD16/32^+^; GMP) numbers in P0, P14 or 8 weeks old adult mice (Figure 3L-T; Figure S1Q, R; Figure S2).^27,28^ Control and *Skida1^Δ/Δ^* progenitors had similar colony forming unit (CFU) frequencies in methylcellulose media supplemented with myeloid and erythroid cytokines (Figure 3U, V). Altogether, the data show that *Skida1* has little to no role in normal fetal, neonatal or adult hematopoiesis.

### *Skida1* selectively sustains neonatal MLL::ENL-expressing HSCs and MPPs

We next tested whether *Skida1* regulates hematopoiesis and leukemogenesis in the context of MLL::ENL expression. We generated mice with *Skida1^flox^* and previously described Tet-Off-ME (*Vav1-Cre; Rosa26^LoxP-STOP-LoxP-tet-transactivator^*; *Col1a1^TetO-MLL::ENL^*) alleles and evaluated HSC and MPP3/4 numbers at P0, P14 and P28. The Tet-Off-ME system express MLL::ENL in developing blood progenitors in the absence of DOX, beginning at E10.5.^10^ *Skida1* deletion caused no significant changes in MLL::ENL-expressing HSC or MPP3/4 numbers at P0 (Figure 4A, B). However, by P14, we observed severe depletion of HSCs, and to a lesser extent MPP3/4s, in *Skida1^Δ/Δ^;* Tet-Off-ME mice (Figure 4C, D). P28 *Skida1^Δ/Δ^;* Tet-Off-ME mice showed sustained HSC depletion, but MPP3/4 numbers recovered and were similar to *Skida1^+/+^; Tet-Off-ME* controls (Figure 4E, F). Thus, *Skida1* helps maintain the HSC/MPP population in MLL::ENL-expressing mice prior to transformation, but progenitor depletion does not occur until shortly after birth, and MPP depletion is only transient. The data indicate a temporally restricted role for *Skida1* in pre-leukemic hematopoiesis.

**Figure 4.**
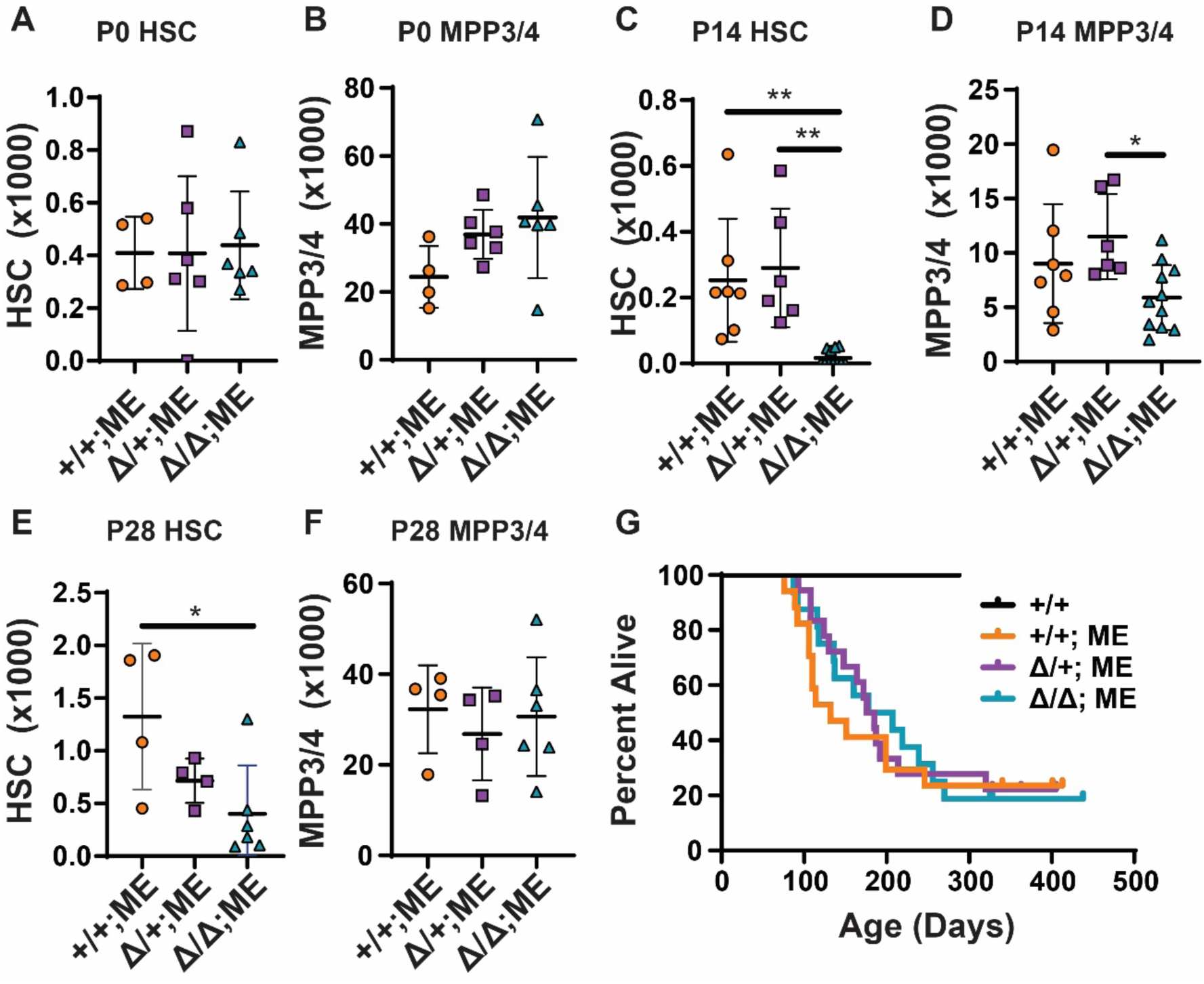
*Skida1* sustains neonatal MLL::ENL-expressing HSCs and MPPs. (A-F) HSC and MPP3/4 numbers for the indicated genotypes in P0 whole liver (A, B), P14 bone marrow (C, D), or P28 bone marrow (E, F). Panels C-F indicate cell numbers for two hindlimbs (tibia plus femur). P0 n= 4 (+/+;ME), n=6 (Δ/+;ME), n=6 (Δ/Δ;ME); P14 n=7 (+/+;ME), n=6 (Δ/+;ME), n=11 (Δ/Δ;ME); P28 n=4 (+/+;ME), n=4 (Δ/+;ME), n=6 (Δ/Δ;ME). *p<0.05, **p<0.01 by one-way ANOVA with Tukey’s posthoc test. (G) Kaplan-Meier survival curves for Tet-Off-ME mice of the indicated *Skida1* genotypes. n= 6 (+/+), n= 14 (+/+;ME), n= 14 (Δ/+;ME), n= 13 (Δ/Δ;ME).

We next evaluated longitudinal survival of Tef-Off-ME mice after deleting one or both *Skida1* alleles. Some, but not all, fully transformed Tet-Off-ME AML express *Skida1* (Figure S3A). We therefore anticipated a delay but not a complete abrogation of leukemogenesis, given that HSC/MPP depletion in *Skida1^Δ/Δ^;* Tet-Off-ME neonates is transient and not all AML express the *Skida1* gene. Instead, *Skida1* deletion had no effect on survival (Figure 4G). All mice died of morphologically and phenotypically similar AML, irrespective of *Skida1* genotype (Figure S3B). Thus, despite having a role in maintaining neonatal MLL::ENL-expressing HSC/MPPs prior to the onset of AML, *Skida1* is not required for myeloid leukemogenesis.

### *Skida1* deletion causes loss of B-cell identity programs and B-cell potential in neonatal MLL::ENL-expressing HSCs

One explanation for why leukemogenesis proceeds normally in *Skida1^Δ/Δ^;* Tet-Off-ME mice, despite neonatal HSC/MPP depletion, is that the Tet-Off-ME line develops AML exclusively, and *Skida1* might selectively maintain lymphoid-biased, MLL::ENL-expressing progenitors. To assess B-cell priming and other population-specific gene expression changes, we performed Cellular Indexing of Transcriptomes and Epitopes (CITE-seq) on P0 control (*Vav1-Cre* negative), *Skida1^+/+^;* Tet-Off-ME and *Skida1^Δ/Δ^;* Tet-Off-ME Lineage^-^c-Kit^+^ (LK) progenitors. We performed the assays at P0 because this timepoint preceded the depletion of progenitors observed at P14, and it allowed us to capture HSCs in the dataset. The data were analyzed by Iterative Clustering with Guide Gene Selection, version 2 (ICGS2)^29^ to generate clusters and visualize the data by Uniform Manifold Approximation and Projection (UMAP) (Figure 5A). We annotated the clusters based on antigen derived tags and gene expression as previously described^30,31^ and identified populations of phenotypic HSC, MPP3/4 and GMPs (Figure 5B, Figure S4). *Skida1^+/+^;* Tet-Off-ME and *Skida1^Δ/Δ^;* Tet-Off-ME progenitors had similar distributions across the UMAP and similar frequencies of each cell population (Figure 5B, C). MLL::ENL caused MPP3/4 cluster expansion, at the expense of HSCs and GMPs, irrespective of *Skida1* genotype. To identify *Skida1*-dependent changes in gene expression, we generated pseudobulk populations and compared gene expression in *Skida1^+/+^;* Tet-Off-ME and *Skida1^Δ/Δ^;* Tet-Off-ME HSC- and MPP3/4-enriched clusters (clusters 12 and 23, respectively; Tables S5 and S6).^32^ GSEA with Hallmark gene sets revealed that *Skida1* deletion resulted in modest but significant enrichment of genes associated with cell cycle and metabolism (e.g., E2F targets, MYC targets, and G2M checkpoint signatures) (Figure 5D).^33^ A B-cell signature was significantly enriched in *Skida1^+/+^;* Tet-Off-ME HSCs and MPPs, relative to *Skida1^Δ/Δ^;* Tet-Off-ME HSCs and MPPs (Figure 5E). This observation suggests that *Skida1* deletion reduces B-cell priming in newborn, MLL::ENL-expressing HSCs and MPPs that ultimately manifests as a transient reduction in numbers.

**Figure 5.**
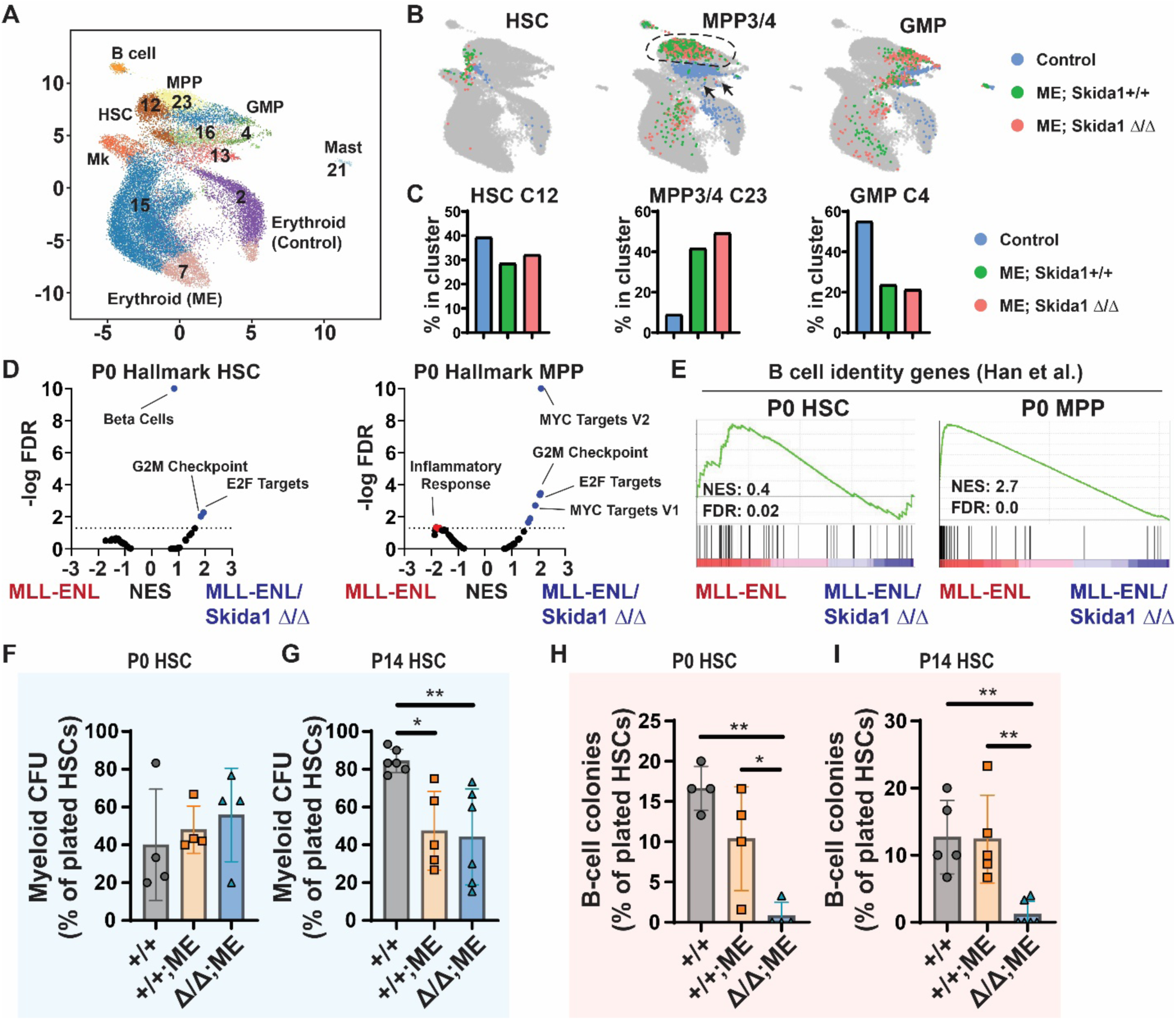
*Skida1* promotes B-cell priming and supports B-cell potential in MLL::ENL-expressing neonatal progenitors. (A, B) UMAPs representing single cell gene expression in P0 LK cells from control (*Vav1-Cre* negative), *Skida1*^+/+^; Tet-Off-ME and *Skida1*^μ/M^; Tet-Off-ME mice. Cluster identities were assigned by ICGS2 and are indicated. Cells were annotated based on antigen derived tags and gene expression as shown in Figure S4. (C) Distribution of cells for each of the indicated genotypes that map to clusters 12 (HSC-enriched), cluster 23 (MPP3/4-enriched), and cluster 4 (GMP-enriched). (D) Volcano plots showing enrichment of Hallmark gene sets within the indicated genotypes as measured by GSEA performed to compare *Skida1*^+/+^; Tet-Off-ME and *Skida1^Δ/Δ^;* Tet-Off-ME HSCs (cluster 12) and MPP3/4s (cluster 23). n=4 pseudoreplicates. (E) GSEA plot showing reduction of B-cell identify genes in P0 HSC and MPP clusters in the absence of *Skida1*. (F, G) Myeloid CFU frequencies from P0 liver (F) and P14 bone marrow (G) HSCs. P0 n= 4 (+/+), n=4 (+/+;ME), n=4 (Δ/Δ;ME); P14 n=6 (+/+), n=5 (+/+;ME), n=6 (Δ/Δ;ME). (H, I) B-cell colony formation frequencies from P0 liver (F) and P14 bone marrow (G) HSCs. P0 n= 4 (+/+), n=4 (+/+;ME), n=4 (Δ/Δ;ME); P14 n=5 (+/+), n=5 (+/+;ME), n=6 (Δ/Δ;ME). *p<0.05 and **p<0.01 by one-way ANOVA with Tukey’s posthoc test.

We tested whether *Skida1* deletion alters the myeloid and B-cell potential of MLL::ENL-expressing HSCs by performing myeloid and B-cell colony forming assays. Myeloid colony formation was assessed by sorting single P0 or P14 HSCs into methylcellulose media that contained myeloid and erythroid cytokines and then counting colonies 14 days later. B-cell potential was assessed by sorting single HSCs into media supported by IL-7, FLT3-L, and OP9 stomal cells.^30^ We then tested for B-cell production 21 days later by flow cytometry (Figure S5A, B). MLL::ENL had no effect on myeloid colony formation at P0, though it led to a modest reduction in myeloid CFU at P14 (Figure 5F, G). *Skida1* deletion had no effect on myeloid CFU at either age (Figure 5F, G). In contrast, *Skida1* deletion led to near-complete loss of HSC B-cell potential at both P0 and P14 (Figure 5H, I). Of note, we did not see a reduction in HSC-derived B-cell colonies when *Skida1* was deleted on a non-MLL::ENL-expressing background (Figure S5C). The data show that *Skida1* regulates B-cell priming and potential in neonatal MLL::ENL-expressing HSCs.

### SKIDA1 is not required to sustain fully-transformed infant B-ALL

Mouse models pose a challenge for investigating the evolution of MLLr infant B- ALL because MLL fusion proteins tend to drive AML rather than B-ALL in mice.^10,13,34–37^ Even when B-ALL does arise (as in the context of MLL::AF4 expression), the latency far exceeds the neonatal developmental window.^38,39^ We therefore used patient-derived xenograft models (PDXs), generated from infants with either MLL::ENL or MLL::AF4 leukemias, to test whether SKIDA1 is necessary to sustain infant B-ALL.^40,41^ We used CRISPR/Cas9 to create targeted insertion-deletion mutations within *SKIDA1* (targeting the N-terminal region of the protein) or a non-coding control site within the *AAVS1* gene (Figure 6A). Edited cells were then transplanted into immunodeficient NOD.Cg-*Prkdc^scid^ Il2rg^tm1Wjl^*/SzJ (NSG) mice. We monitored recipient survival, recognizing that differences might not be evident due to rapid growth of unedited cells. Indeed, across 5 independent PDX models, we did not observe survival differences between recipients of *AAVS1-* and *SKIDA1-*edited cells (Figure 6B and data not shown).

**Figure 6.**
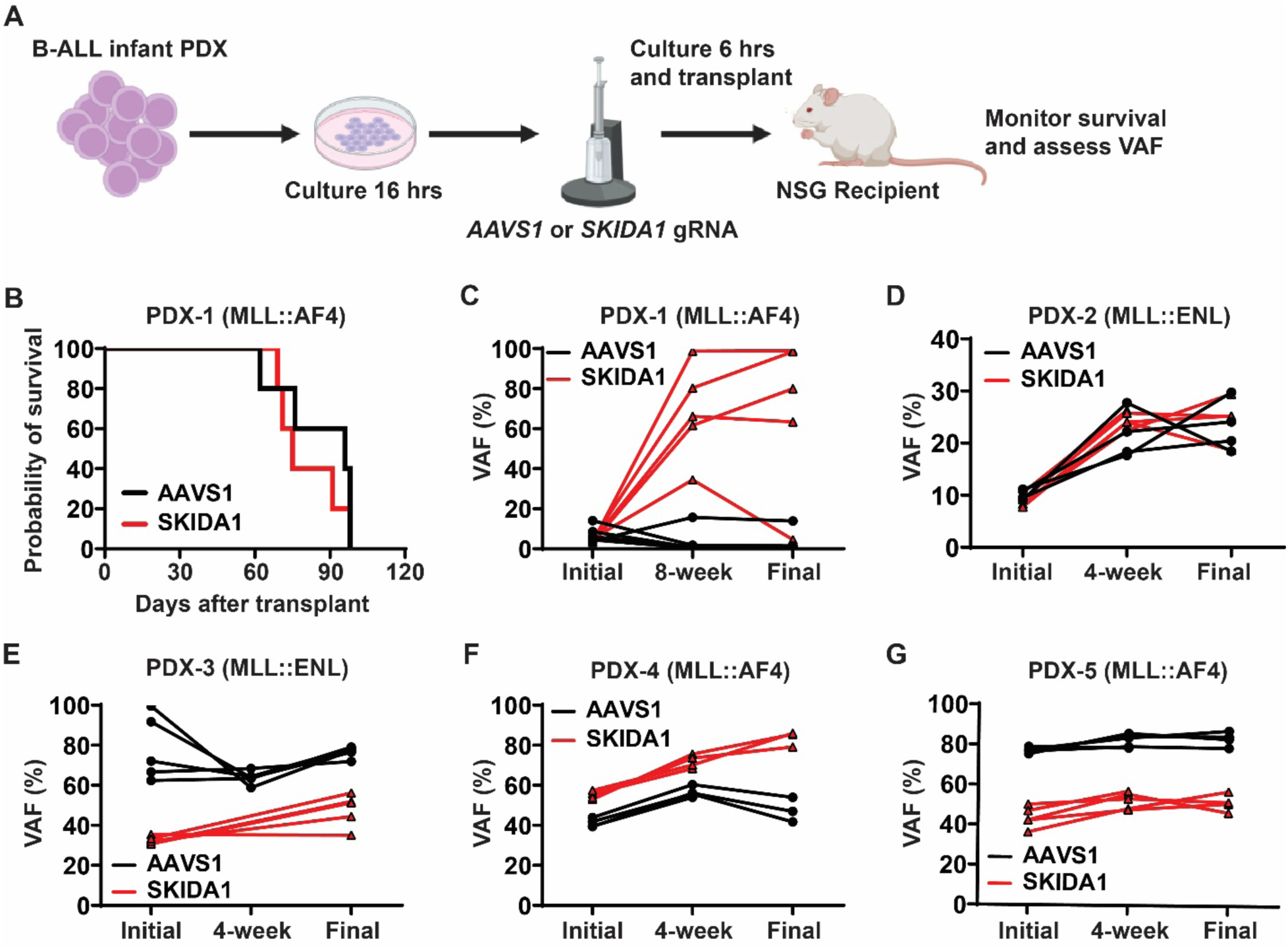
*SKIDA1* is not required to sustain infant B-ALL. (A) Experimental design for CRISPR/Cas9-mediated inactivation of *SKIDA1* in infant B- ALL PDX. (B) Kaplan-Meier survival curves for NSG recipients of AAVS- or SKIDA1- edited infant B-ALL PDX-1. (C-G) Initial and post-transplant VAF levels in cultured PDX cells (initial, collected at 72 hours after editing), or in engrafted cells at the indicated timepoints and at time of death (final, 8-12 weeks). Five unique PDX were evaluated, and the underlying driver mutation is indicated for each (MLL::AF4 or MLL::ENL). PDX-1 n= 5 (*AAVS1*) and n=5 (*SKIDA1*); PDX-2 n= 4 (*AAVS1*) and n=5 (*SKIDA1*); PDX-3 n= 5 (*AAVS1*) and n=5 (*SKIDA1*); PDX-4 n= 4 (*AAVS1*) and n=4 (*SKIDA1*); PDX-5 n= 5 (*AAVS1*) and n=5 (*SKIDA1*).

In addition to survival monitoring, we tracked variant allele frequencies (VAF) in the *AAVS* or *SKIDA1* genes prior to transplant (initial), after engraftment (4- or 8-weeks after transplant), and at the time of death. In the post-transplant periods, human B-ALL cells were isolated by flow cytometry prior to assessing VAF. If *SKIDA1* is necessary to sustain infant B-ALL, then *SKIDA1* VAF should have declined over time in edited leukemias, even if survival itself was unaffected due to growth of unedited cells. Instead, recipients of one PDX model showed expansion of *SKIDA1*-mutant cells over time, while the other four PDX models showed relatively stable VAF over time, with almost all variants reflecting out-of-frame insertion-deletion mutations (Figure 6C-G, Figure S6). Altogether, the data do not support a scenario in which fully transformed infant B-ALL cells require *SKIDA1* to sustain the malignancy. The murine data argue for a role for *SKIDA1* in promoting B-cell identity in neonatal MLLr HSCs and MPPs, and for sustaining these progenitors during neonatal stages of life, but *SKIDA1* does not appear to be necessary after transformation.

## DISCUSSION

Mechanisms that define lineage bias in MLLr leukemias remain poorly understood. While MLL fusion proteins drive both B-ALL and AML, their prevalence within each leukemia subtype varies with age. For example, B-ALL accounts for 70-80% of all MLLr leukemia diagnoses in children <1 year old.^1,42–44^ At later ages, the incidence MLLr AML exceeds that of infant ALL.^44,45^ The data suggest that in humans, infancy favors B-cell leukemogenesis. This has been difficult to model in mice due to the strong predilection towards AML, but some mechanisms have been identified. For example, CCL5 expression in adult bone marrow inhibits B-cell leukemogenesis.^36^ Likewise, fetal-specific Igf2/Igf2r and Fn1/VLA-4 may promote lymphoid bias.^46^ Mechanisms underlying the lymphoid bias of fetal/neonatal MLLr progenitors are therefore captured in mouse models, even if the models ultimately develop AML.

Here we identified *Skida1* as an effector of fetal/neonatal B-cell priming, specifically in the context of MLL fusion protein expression. SKIDA1 associates with PRC2 yet it has no clear role in normal hematopoiesis. This lack of phenotype in *Skida1* deficient mice implies that SKIDA1 has only a limited effect on PRC2 function overall because other PRC2 proteins, such as EED and either EZH1 or EZH2, are required for fetal, neonatal and adult HSC function.^23,26^ SKIDA1 helps sustain MLL::ENL-expressing HSCs and MPPs through neonatal and juvenile stages of life. Interestingly, this dependence is temporally restricted, as it does not arise until shortly after birth, and MLL::ENL-expressing MPP numbers recover by P28. SKIDA1 is necessary to support B-cell potential in newborn and neonatal MLL::ENL-expressing HSCs, and it may therefore contribute to infant B-ALL evolution rather than B-ALL maintenance. SKIDA1 appears to have little to no role in myelopoiesis or myeloid leukemogenesis, unlike other PRC2 proteins.^21,22^ Thus, SKIDA1 activity shapes some but not all of the cellular and molecular consequences of MLL fusion protein expression.

The data illustrate how select, temporally-restricted targets of MLL fusion proteins can tune cell fates at different ages. Prior studies have shown that HSCs and MPPs begin acquiring adult-like properties during late gestation, and the transition to adult identity then proceeds gradually through neonatal and juvenile stages of life.^47,48^ While MLLr often originate fetal HSC/MPPs, the data implicate neonatal programs as critical determinants of SKIDA1 dependence and lymphoid bias.

## METHODS

### Mouse lines and patient-derived xenograft models

*Skida1* germline loss-of-function mice were generated by deleting the single *Skida1* exon with CRISPR/Cas9. Blastocysts were electroporated with gRNAs complexed with Cas9 to target regions 5’ (CCAGGTACTGATCCGCATCGNGG) and 3’ (TGTAACACCATTAATGGACTNGG) to the coding exon. Deletion was confirmed by next generation sequencing (NGS). *Skida1* floxed alleles were generated with the same Cas9/gRNA complexes with single strand oligonucleotides co-electroporated to insert LoxP sites. LoxP insertion was confirmed to be in cis by long range PCR and NGS. *Vav1-Cre*, *Rosa26^LSL-rtTA-IRES-mKate^*^2^, and *Rosa26^LSL-tTA^* were obtained from Jackson Laboratory.^49–51^ *Col1a1^TetO-MLL::ENL^*mice (from David Bryder, Lund) were described previously.^13^ All strains are on a C57BL/6 genetic background. DOX chow (200 ppm) was purchased from Bioserv. Infant B-ALL PDX were generated from clinically annotated primary specimens, without protected health information, collected under the auspices of IRB-approved research protocols at Washington University or Children’s Hospital of Philadelphia, as previously described.^40,41^ All procedures were performed according to Institutional Animal Care and Use Committee-approved protocols.

### Flow cytometry and transplantations

Cells were isolated, stained, analyzed, and transplanted as previously described.^10,47,52^ All antibodies are from Biolegend.

### RNA-seq

Fifty thousand LSK cells were isolated from Tet-On-ME mice at the indicated ages by flow cytometry and cultured with and without DOX for 48 hours. The cells were then washed, pelleted by centrifugation and resuspended in RLT-plus RNA lysis buffer (QIAGEN). RNA-seq libraries were generated an analyzed as previously described.^31,47^

### CITE-seq library construction and analysis

CITE-seq libraries were generated as previously described.^30,31^ The Cell Ranger v6.1.2 pipeline was used to align, filter and normalize digital gene expression files as previously described.^30,31^ We performed Iterative Clustering and Guide-gene Selection (ICGS, version 2) analysis with the AltAnalyze toolkit^29^ to visualize cells and annotate clusters. For differential expression analyses, Presto was used to generate pseudoreplicates within clusters (4 per sample group) by aggregating read counts within each replicate.^32^ DESeq2 was then used to identify differentially expressed genes.^53^ Pseudoreplicates were used to perform to perform GSEA.^33^

### Immunoprecipitation and mass spectrometry

Cell lines were cultured for three days with DOX (2ug/mL) to induce SKIDA1- FLAG. On day three, ten million cells were lysed in mammalian cell lysis buffer (50mM Tris-HCl + 150mM NaCl + 0.1% NP-40 + protease inhibitor) and supernatant was incubated with anti-FLAG M2 magnetic beads (Sigma, M8823) overnight. The beads were then washed with MCLB and on-bead protein digestion by trypsin was performed, followed by liquid chromatography – mass spectrometry (LC-MS/MS). Scaffold 5 software was used to analyze mass spectrometry data. To identify potential binding partners, the data were filtered using a protein probability of 95%, a minimum of two unique peptides identified, and 90% peptide probability (spectral counts are shown in Table S2 and S3).

### Western blots

Western blots were performed as previously described.^52^ Antibodies include the following: FLAG M2 (Sigma, F1804) alpha-Tubulin (Cell Signaling, 2144S), EZH2 (Cell Signaling, #5246), SUZ12 (Cell Signaling, #3737), and MTF2 (Proteintech, #16208-1- AP).

### Myeloid and B-cell potential assays

Myeloid and B-cell colony formation assays were performed as previously described.^30^

### CRISPR/Cas9 editing and transplantations

B-ALL PDXs were thawed, washed with 40% FBS + 60% PBS, and cultured overnight in RPMI + Flt3-L (25ng/mL) + IL-7 (10ng/mL) + 1% P/S prior to electroporation. Individual electroporation reactions were performed for each recipient mouse. Ribonucleoprotein (RNP) complex was prepared by incubating Cas9 (IDT) with either *SKIDA1* gRNA (GACGACCGTGCACAAGCGCA) or a control *AAVS1* gRNA (CCAGTGTGCCGCTGCGTGCA) at a 6:1 molar ratio at room temperature for 10 minutes. 100-200 thousand cells were electroporated with RNP (1600V/ 10ms/ 3 pulses) and cultured for 6 hours. Electroporated cells were then transplanted into NSG mice by retro-orbital injections. VAF were obtained by preforming next generation sequencing and analyzing the data using CRISPResso2.^54^

## DATA AVAILABILITY

CITE-seq and RNA-seq data have been deposited in Gene Expression Omnibus.

Accession numbers GSE239107 and GSE239108.

## AUTHOR CONTRIBUTIONS

J.A.M. designed and oversaw all experiments, interpreted the data and wrote the manuscript with J.M.-C. J.M.-C. designed, conducted, and interpreted the experiments.

W.Y. performed all bioinformatic analyses. E.D., R.M., E.B.C., and R.M.P. performed the experiments. S.K.T. provided infant leukemia PDX specimens. All authors reviewed the manuscript.

## ACKNOWLEDGMENTS

We thank David Bryder for providing TetO-MLL::ENL mice. We thank J. Michael White for assistance in generating the *Skida1* loss-of-function mice. This work was supported by grants to J.A.M. from the NHLBI (R01HL152180), the NCI (R01CA285272), Gabriel’s Angel Foundation and the Children’s Discovery Institute of Washington University and St. Louis Children’s Hospital. J.M.-C. was supported by a NCI career development grant (F31CA268923). S.K.T. was supported by NCI U01CA232486 and U01CA243072 awards, a Pennsylvania Department of Health Commonwealth Universal Research Enhancement Program CURE award, and the SchylerStrong Foundation. S.K.T. and J.A.M. are Scholars of the Leukemia and Lymphoma Society.

