## Supplementary Materials for "SKIDA1 transiently sustains MLL::ENL-Expressing hematopoietic progenitors during neonatal stages and promotes B-lineage priming"

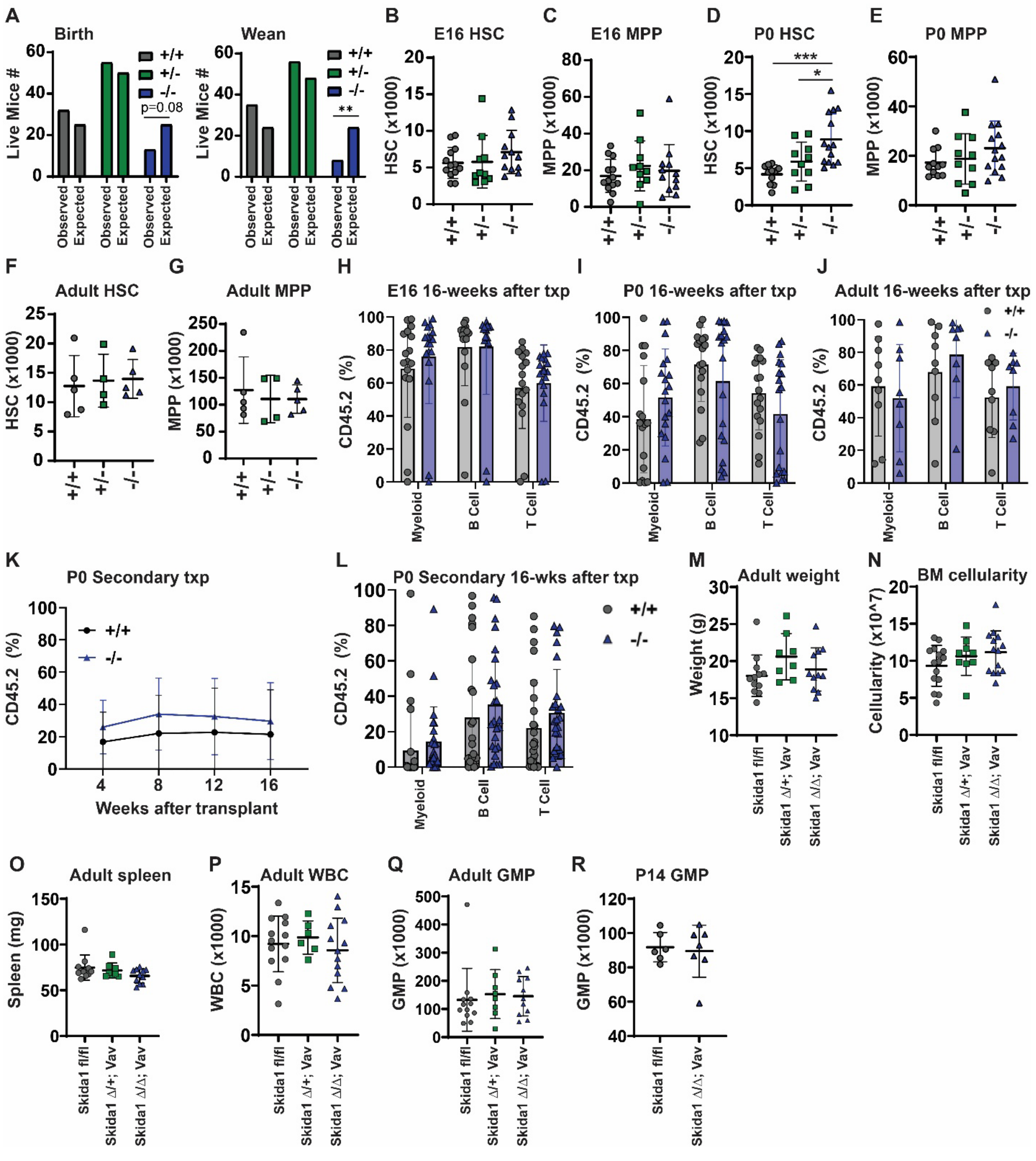


**Supplementary Figure 1 (related to Figure 3).** ***Skida1*** **is not required for normal hematopoiesis.**

(A) Bar graph comparing observed and expected numbers of mice of the indicated genotypes at birth and P21 (weaning age), based on crosses of *Skida1*$^{+/-}$ mice and anticipate Mendelian allele distributions. **p<0.01 by Chi squared test. (B-G) HSC and MPP for mice with indicated *Skida1* genotypes at E16, P0 and 8-10 weeks old (adult). E16 n=13 (+/+), n=10 (+/-), and n=12 (-/-); P0 n=11 (+/+), n=10 (+/-), and n=14 (-/-);

Adult n=5 (+/+), n=4 (+/-), and n=5 (-/-). *p<0.05 and ***p<0.001. (H-J) Donor (CD45.2) lineage-specific chimerism in peripheral blood of recipient mice at 16 weeks after transplantation. Donor ages are indicated. E16 recipients n=17 (+/+) and n=18 (-/-); P0 recipients n=17 (+/+) and n=19 (-/-); Adult recipients n=9 (+/+) and n=8 (-/-). (K) Peripheral blood chimerism in secondary recipients of bone marrow derived from P0 *Skida1*$^{+/+}$ or *Skida1*$^{-/-}$ mice. (L) Donor lineage-specific chimerism in recipients from panel K, at 16 weeks post-transplant. P0 secondary recipients n=25 (+/+) and n=25 (-/-) (M-Q) Body weight (M), bone marrow cellularity (N), spleen weight (O), peripheral white blood cell counts (P) and GMP numbers (Q) in adult control or *Skida1*$^{\Delta/\Delta}$ mice. n=15 (+/+), n=9 (Δ/+), and n=14 (Δ/Δ). (R) GMP numbers in P14 in adult control or *Skida1*$^{\Delta/\Delta}$ mice. n=6 (fl/fl) and n=7 (Δ/Δ). None of the comparisons in panels H-R indicated significant differences among the genotypes, as calculated by one-way ANOVA with Tukey's posthoc test.

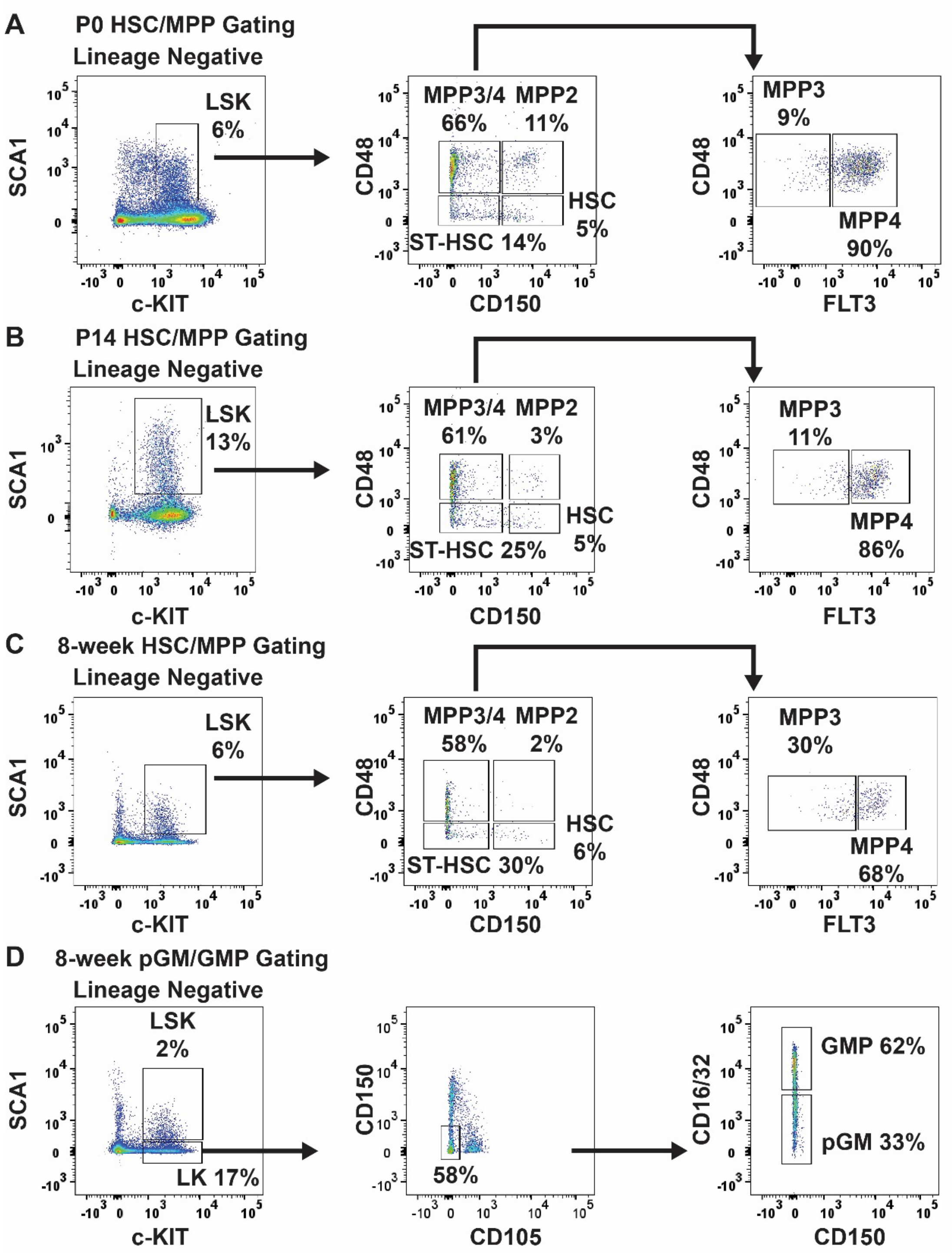


**Supplementary Figure 2 (related to Figure 3). Representative flow cytometry plots and gating strategies for analyzing hematopoiesis in *Skida1* germline- and conditional loss-of-function mice.**

(A-C) Representative flow cytometry plots and gating strategy for HSCs and MPP subpopulations (MPP2, MPP3 and MPP4) at P0, P14 and 8 weeks old, as described by Pietras et al. (Ref. 12). D) Representative flow cytometry plots and gating strategy for committed myeloid progenitors (pGMs and GMPs), as described by Pronk et al. (Ref. 43).

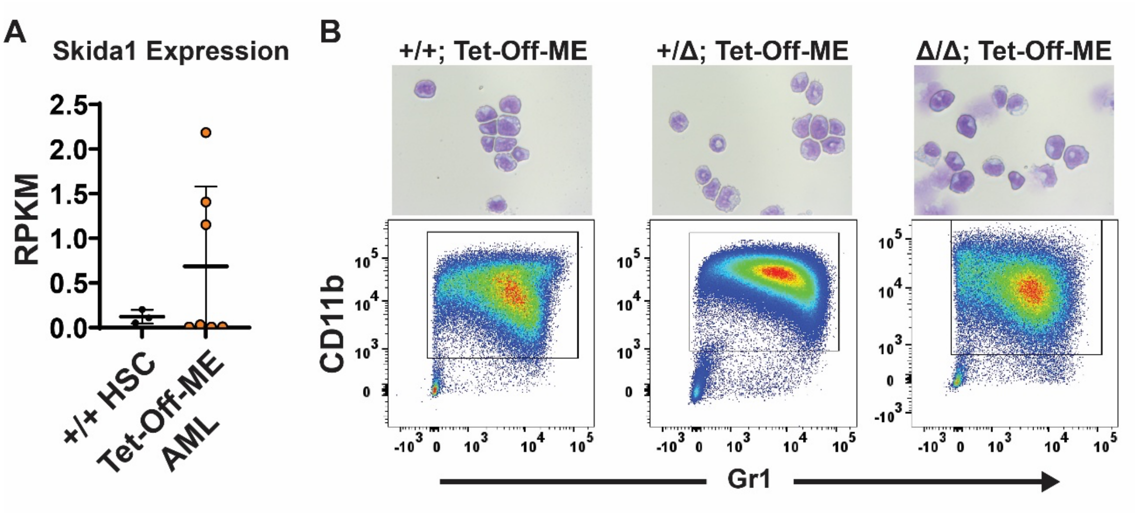


**Supplementary Figure 3 (related to Figure 4). SKIDA1 is not required for myeloid leukemogenesis.**

(A) *Skida1* expression by RNA-seq in wildtype HSCs (n=3) and Tet-Off-ME AML cells (n=7). (B) Representative cytospins showing blast morphologies (top panels) and flow cytometry plots showing myeloid differentiation (bottom panels) in AML from mice of the indicated genotypes.

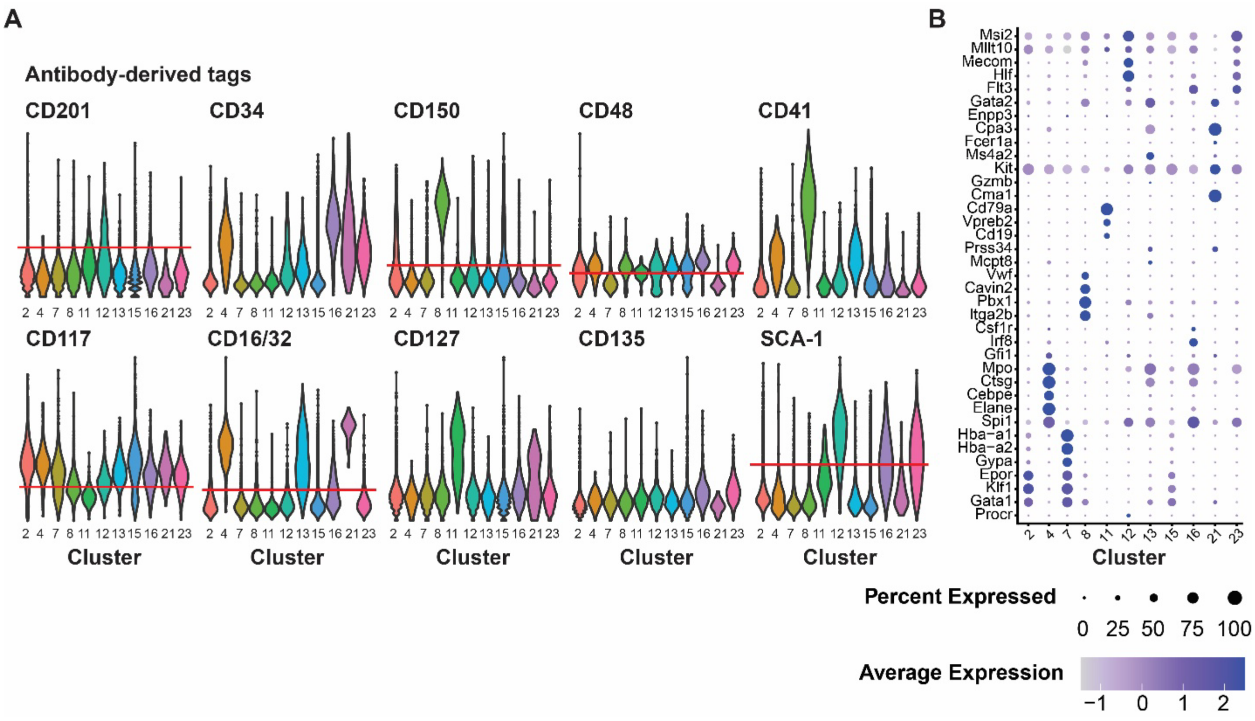


**Supplementary Figure 4 (related to Figure 5). Annotation data for P0 CITE-seq assays.**

(A) Expression of indicated surface markers based on antibody derived tags. Expression levels are shown for each cluster. CD150, CD48, SCA1, CD16-31 and KIT were used to annotate HSC/MPP and pGM/GMP populations. Thresholds for positive versus negative expression are marked by the red line. (B) Expression of lineage-specific marker genes by cluster.

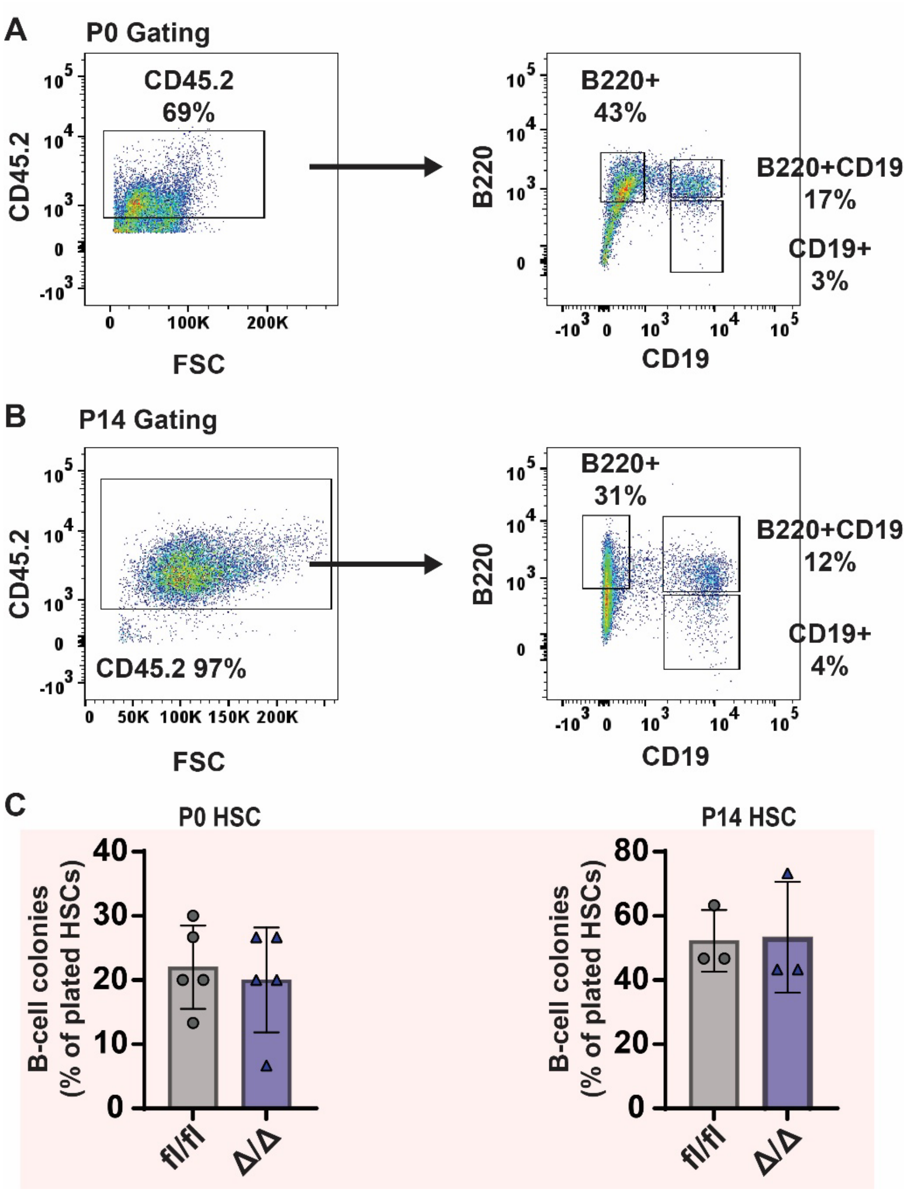


**Supplementary Figure 5 (related to Figure 5). Flow cytometry gating strategy for B-cell colony formation assays.** (A, B) Representative flow cytometry plots and gating strategies to identify CD19 and B220 single- and double-positive B-cells in cultures derived from P0 and P14 HSCs. (C) B-cell colony frequencies from control and Skida1$^{\Delta/\Delta}$ HSCs at P0 (n=5) and P14 (n=3).

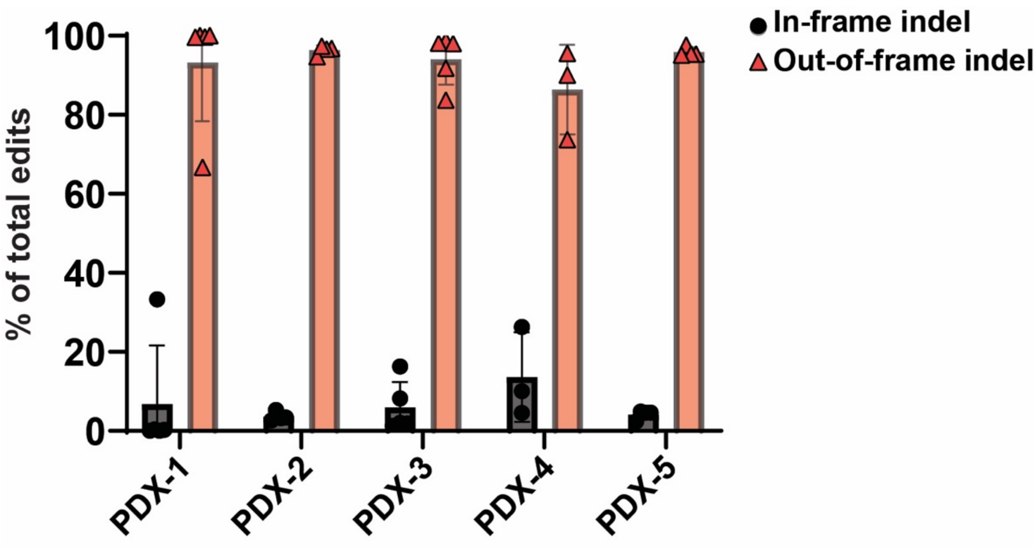


**Supplementary Figure 6 (related to Figure 6). Variant types in Cas9 edited PDX.** The percentage of variants reflecting in-frame and out-of-frame in insertion-deletion mutations for each PDX model. Each data point reflects an independent recipient mouse.
